# Endonuclease G governs an antiviral RNA surveillance system in budding yeast

**DOI:** 10.64898/2026.10.28.762607

**Authors:** Jie Gao, Brittney Remnant, Braden Martyn, Minyoung Chung, Jing Li, Megan Pickett, Sabrina Chau, Mark A. Bayfield, Marc D. Meneghini

## Abstract

Endonuclease G family DNA/RNA nucleases localize to the mitochondrial intermembrane space and prevent escape of mitochondrial DNA to the cytosol or nucleus. While extramitochondrial Endonuclease G also targets nuclear DNA during cell death, RNA regulatory roles of this family remain enigmatic. Here we discover an RNA surveillance system in *Saccharomyces cerevisiae* governed by Nuc1, the yeast Endonuclease G homolog. This system limits replication of L-A, an RNA mycovirus that endemically infects this organism. Whereas L-A resides in the cytosol, we find that Nuc1’s antiviral function requires its mitochondrial intermembrane space localization. To understand Nuc1’s indirect antiviral mechanism we investigated its genetic interactions with *REX1* and *REX2*, RNA exonucleases that control 3’ end processing of tRNA and rRNA intermediates in the nucleus. We demonstrate that the *nuc1*Δ antiviral defect is massively amplified by *rex1*Δ *rex2*Δ leading to L-A RNA levels that surpass mature rRNAs and overwhelm the cells. *NUC1* concurrently controls a pool of mis-processed tRNA and rRNA species that accumulate in *rex1*Δ *rex2*Δ mutants. Linking these mis-processed RNAs with L-A control, we show they are significantly reduced in L-A infected cells, suggesting that L-A depletes them to enhance its replication. Our findings identify a previously unrecognized RNA surveillance function of Nuc1 that prevents aberrant rRNA and tRNA species from promoting replication of the L-A mycovirus.

## Introduction

Endonuclease G (EndoG) belongs to an ancient family of DNA/RNA nucleases distinguished by a ββα-metal structural core and localization to the mitochondrial intermembrane space. EndoG has deep evolutionary roots, and the phylogenetic distribution of ββα-metal proteins suggests its origin predates the alphaproteobacterial ancestor of mitochondria. Duplication of an ancestral eukaryotic gene occurred sometime after the split of animalia and fungal lineages, giving rise to EndoG and its closely related paralog ExoG in metazoans^1^. In bacteria, NucA and other ββα-metal proteins are secreted as pro-virulence factors that degrade DNA-rich neutrophil extracellular traps (NETs) thereby promoting immune evasion^2,3^. A similar scenario has recently been shown in the fungal plant pathogen *Puccinia striiformis* during which EndoG secretion suppresses plant extracellular DNA-triggered immunity^4^. Evocative of these NucA/EndoG secretion functions, early studies showed that EndoG is released from mitochondria to execute nuclear genome fragmentation during apoptotic death in mouse cells^5^. Within mitochondria, EndoG functions during the elimination of paternal mitochondrial DNA in invertebrates and prevents the escape of mitochondrial DNA to the cytosol in mouse embryonic fibroblasts^6–8^.

Biochemical RNase activity of EndoG was first shown for *Saccharomyces cerevisiae* (yeast) and bovine EndoG^9,10^. Subsequent studies of human EndoG revealed RNA to be the strongly preferred *in vitro* substrate^11^. In mouse cells, EndoG can generate the RNA primers required by DNA polymerase γ to initiate mitochondrial DNA replication^10^. In human hepatic cells, EndoG removes 3′-end extensions of mitochondrial polycistronic RNA transcripts to promote proper folding of mitochondrial-tRNA^Thr^, boosting mitochondrial function^12^. In the fission yeast *Schizosaccharomyces pombe*, deletion of the EndoG homolog *pnu1* slows the rate of bulk RNA decay during nitrogen starvation, suggesting a broad RNase function in this nutritional context^13^. Beyond these studies, little is known of EndoG RNA regulatory functions.

Similarly to its mammalian EndoG homologs, yeast Nuc1 prevents the escape of mitochondrial DNA to the nucleus^14,15^. Remarkably, Nuc1 also targets nuclear DNA during death of oxidatively stressed mitotic cells or of discarded meiotic products during sporulation^16,17^. In addition to these DNA targets, Nuc1 has a unique function in repression of L-A, an RNA virus that infects all lab strains and many environmental isolates^18–20^. L-A belongs to the *Totiviridae* family of double stranded RNA viruses, and like all Totiviruses, L-A replicates in the cytosol and persists as a long-term endemic infection^21^. While extramitochondrial Nuc1 accumulates during sporulation or in oxidatively stressed cells, Nuc1 is only found in mitochondria in healthy mitotic cells where it represses L-A in a manner dependent on its catalytic activity^16,19^. These highlight a central question: how does a mitochondrial nuclease repress a cytosolic RNA virus?

Here we determine the antiviral mechanism of Nuc1 in mitotic cells, confirming that it performs this function in the mitochondrial intermembrane space. To this end, we show that cytosolic Nuc1 is ineffective at L-A repression, highly toxic, and is opposed by a cytosolic Nuc1 binding protein we name Nic1 (<u>N</u>uc1 inhibitor in the <u>c</u>ytosol). To understand *NUC1*’s indirect antiviral mechanism, we investigated its genetic interactions with *REX1* and *REX2*, 3’-5’ RNA exonucleases that control 3’ end processing of nuclear tRNA and rRNA processing intermediates^22–28^. While *rex1*Δ *rex2*Δ double mutants exhibit no antiviral defect, loss of *NUC1* in a *rex1*Δ *rex2*Δ strain causes a massively synergistic >200-fold increase in L-A, leading to viral dsRNA levels that surpass mature 25S and 18S rRNAs. These findings can be explained by a model in which Nuc1 targets aberrant, pro-viral RNA species that accumulate in *rex1*Δ *rex2*Δ cells. Northern blotting experiments with the known Rex1/2 targets 5S rRNA and tRNA-Arg3 support this model and reveal that these species are significantly depleted in L-A infected cells, suggesting that they may be the source of NTPs necessary for the massive level of L-A replication. Our findings reveal a novel RNA surveillance role of Nuc1 that counters aberrant pro-viral RNA species.

## Results

### Nuc1 mitochondrial targeting prevents toxicity caused by its cytosolic accumulation

To determine where Nuc1 functions to repress L-A we engineered alleles that affect its accumulation in the mitochondria. The Nuc1 N-terminal targeting region contains a hydrophobic stretch flanked by positively charged residues, characteristic of the signal-anchor mitochondrial targeting domains found in Tom20, Tom70, and Om45 (Fig. 1a)^29^. While these proteins are anchored to the outer mitochondrial membrane, biochemical studies showed a physical association of Nuc1 with the mitochondrial inner membrane with its nuclease domain facing the intermembrane space, suggesting that signal-anchor domains can also tether proteins to the inner mitochondrial membrane^16^. Indeed, ExoG is also tethered to the inner mitochondrial membrane via an N-terminal hydrophobic region, unlike EndoG that is targeted via a cleavable N-terminal sequence (Fig. 1a)^1^. Nuc1 may therefore be regarded as more ExoG like, and phylogenetic analysis suggests that the solitary EndoG family protein in fungi possesses functions that were split between EndoG and ExoG in animals^1^. We confirmed that N-terminal Nuc1 sequences comprising the hydrophobic stretch only (N24) or a larger N43 fragment were sufficient to localize GFP to the mitochondria (Fig 1a, b).

**Figure 1:**
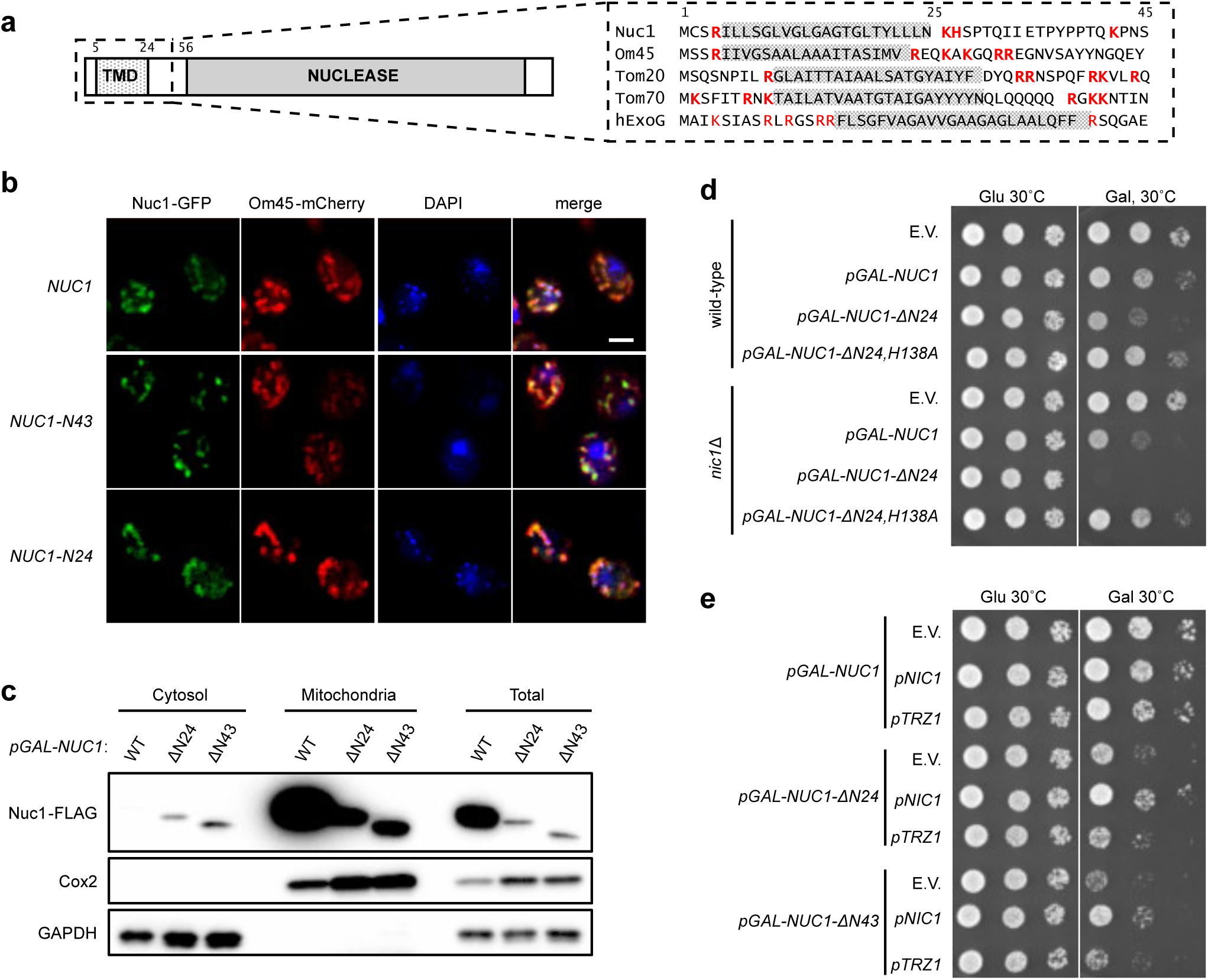
Cytosolic Nuc1 is toxic to the host. **(a)** N-terminal peptide sequence of Nuc1 and other mitochondrial signal-anchor proteins. The hydrophobic stretch of amino acids is shaded in grey while the flanking positively-charged residues is noted in red. **(b)** Fluorescence microscopy of the subcellular localization of Nuc1. Nuc1 or a fragment of its N-terminus was tagged with GFP to monitor its subcellular localization. Mitochondria were marked with the mitochondrial protein OM45-mCherry, and cells were stained with DAPI to visualize nuclei. Scale bar, 1 μm. **(c)** Western blot of Nuc1-FLAG of cell fractions from wild-type strains with plasmid expressing either wild-type *NUC1*, or N-terminus deleted *NUC1* alleles. GAPDH and Cox2 were used to represent the cytosolic and mitochondrial fractions, respectively. The amounts of protein loaded in the mitochondrial and total fractions were 25% of other samples. **(d)** Spot analysis of wild-type and *nic1Δ* strains with plasmid expressing either wild-type *NUC1* or its variants. *NUC1* overexpression was induced by galactose treatment. **(e)** Spot analysis of wild-type strains with plasmid expressing either wild-type *NUC1* or N-terminus deleted *NUC1* alleles along with high-copy plasmids of *NIC1* or *TRZ1*. *NUC1* overexpression was induced by galactose treatment.

To further investigate Nuc1 targeting we constructed plasmids enabling galactose-inducible expression of C-terminal FLAG tagged Nuc1 or versions lacking the N-terminal 24 or 43 amino acids (“Nuc1-Δ24” and “Nuc1-Δ43”). Following two hours of galactose induction, we assessed the presence of Nuc1-FLAG in cytosolic and mitochondrial fractions from cells using western blotting (Fig 1c). Both Nuc1-Δ24 and Nuc1-Δ43 were present at dramatically reduced levels in whole-cell lysates compared with full length Nuc1, suggesting the protein was de-stabilized without these sequences (Fig 1c). While full length Nuc1 was exclusively detected in the mitochondrial fraction, Nuc1-Δ24 and Nuc1-Δ43 were present in the cytosolic fraction, though much of these proteins were still found in the mitochondrial fraction (Fig 1c). These findings confirm a mitochondrial targeting role of the Nuc1 signal-anchor domain but suggest other sequences in Nuc1 may facilitate its association with mitochondria and/or other cellular material in the mitochondrial fraction. When these strains were assessed for growth using spot testing, we found that sustained over-expression of Nuc1-Δ24 or Nuc1-Δ43 was toxic (Fig 1d and S1). Mutation of the key Nuc1 catalytic residue histidine-138 to alanine (Nuc1-H138A) abolished this toxicity, confirming that it was due to rogue nuclease activity (Fig. 1d).

Bacterial NucA is bound by an inhibitory protein to oppose its inherent potential for self-toxicity^30,31^. Accordingly, EndoG is bound by proteins that inhibit its extramitochondrial activity in Drosophila and humans, prompting us to ask whether yeast similarly possess an EndoG-inhibitory protein^32–34^. The *YMR099C* gene encodes a poorly characterized Glucose-6-phosphate 1-epimerase that localizes to the cytosol^35–37^. Numerous high-throughput mass spectrometry and focused co-immunoprecipitation studies identified Ymr099c and Nuc1 in a complex^38–41^. Subsequent *in vitro* experiments showed that Ymr099c forms a heterotetramer with Nuc1 that strongly inhibits its activity^37^. To test if *YMR099C* opposes cytosolic Nuc1 toxicity *in vivo* we expressed the galactose-inducible *NUC1* plasmids in *ymr099c*Δ mutants and found that the toxicities of full length Nuc1, Nuc1-Δ24, and Nuc1-Δ43 were strongly enhanced (Fig 1d and S1). Using high-copy plasmids we further showed that *YMR099C* overexpression rescued the toxicity of Nuc1-Δ24 and Nuc1-Δ43 (Fig. 1e). In contrast, overexpression of *TRZ1*, which encodes a Nuc1-inhibitory protein localized to the mitochondrial matrix and nucleus, failed to rescue toxicity caused by either mutant (Fig. 1e)^37^. These findings support the conclusion that Ymr099c inhibits rogue nuclease activity of cytosolic Nuc1, and we therefore name the encoding gene *NIC1*: <u>N</u>uc1 <u>I</u>nhibitor in the <u>C</u>ytosol-1.

### Nuc1 functions in the mitochondrial intermembrane space to repress L-A replication

We leveraged the above findings to investigate where Nuc1 functions to repress L-A. The toxicity caused by galactose overexpression of Nuc1-Δ24 and Nuc1-Δ43 is confounding for investigating Nuc1 antiviral function. We therefore engineered low-copy plasmids that drive *NUC1* expression using its native promoter for subsequent studies.

To repress L-A replication *NUC1* acts in parallel with the Ski-complex, a conserved translation-coupled mRNA surveillance system that controls 3’-5’ mRNA decay^21^. Double mutants combining *nuc1*Δ and *ski3*Δ exhibit a synergistic increase in L-A levels, and this high viral load causes lethality at elevated temperatures due to overwhelming proteotoxic stress, providing a facile growth assay for Nuc1 antiviral function^19,20^. As expected, expression of full length Nuc1-FLAG dramatically reduced levels of the L-A capsid protein (Gag) in a *nuc1*Δ *ski3*Δ strain and rescued its growth defect at high temperature (Fig 2a, b). Compared with full-length Nuc1, Nuc1-Δ24 and Nuc1-Δ43 were substantially less abundant and less effective at reducing Gag levels or rescuing the growth defect of *nuc1*Δ *ski3*Δ mutants (Fig 2a, b). Quantification of L-A RNA levels using quantitative PCR of reverse transcribed RNA confirmed that Gag levels correlated with L-A copy number (Fig 2c). These results show that Nuc1 mitochondrial targeting sequences are required for its antiviral activity and suggest that its cytosolic form is ineffective for L-A repression.

**Figure 2:**
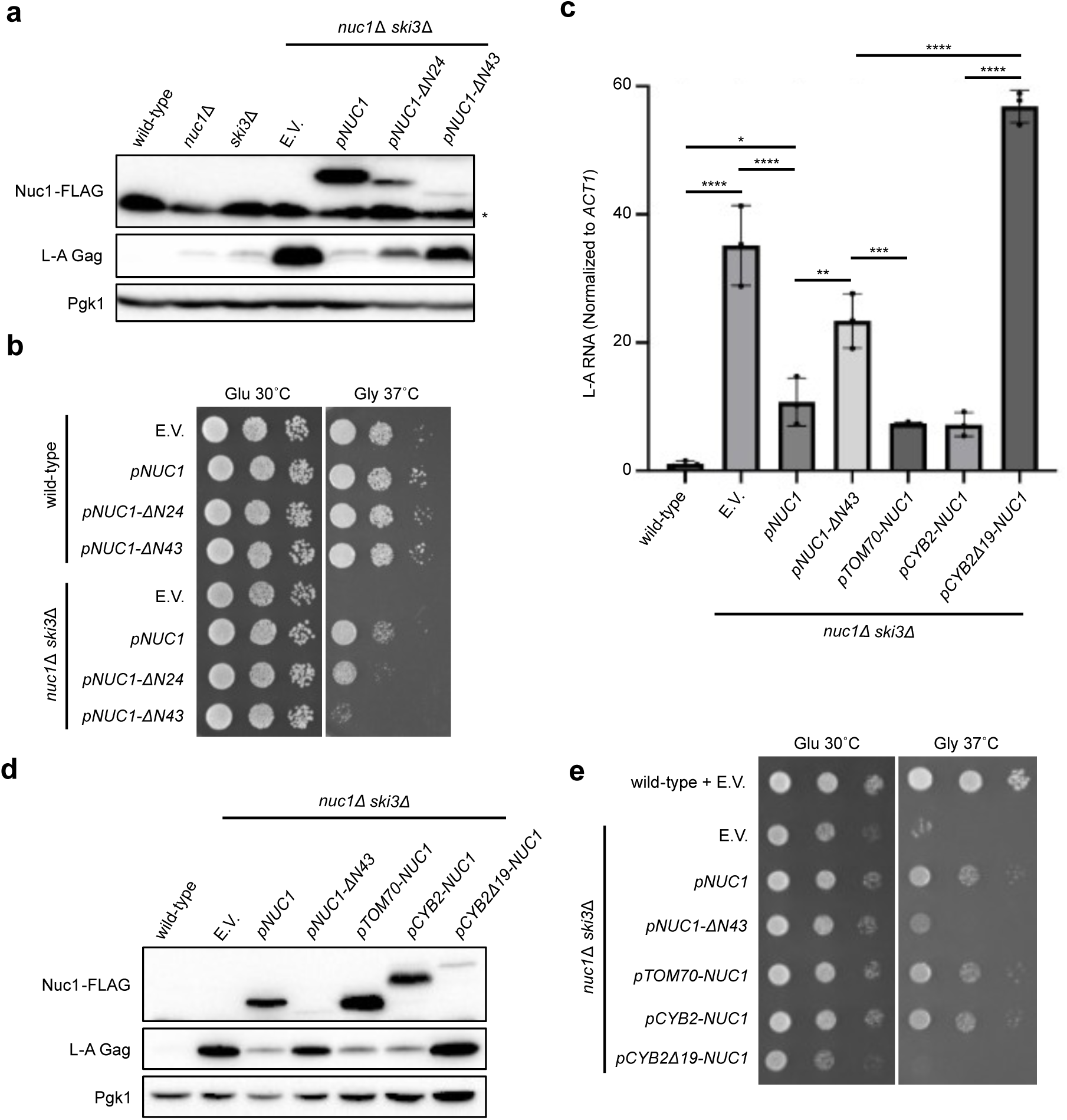
Nuc1 antiviral activity is dependent on its mitochondrial localization. **(a)** Western blot of L-A Gag levels in indicated strains. Nuc1 and its variants were tagged with FLAG epitope tag to confirm expression from plasmid. Pgk1 was used as loading control. **(b)** Spot analysis of wild-type and *nuc1Δ ski3Δ* strains with plasmid expressing either wild-type *NUC1* or N-terminus deleted *NUC1* alleles. Strains were grown at the optimal conditions (glucose at 30°C) and the suboptimal conditions (glycerol at 37°C). **(c)** RT-qPCR quantification of L-A RNA normalized to endogenous *ACT1* RNA in indicated strains. Mean RNA level and standard deviation is shown. n = 3. * p < 0.05, ** p < 0.01, *** p < 0.001, **** p < 0.0001. The p value was calculated using unpaired student’s t-test. **(d)** Western blot of L-A Gag levels in indicated strains. Nuc1 and its variants were tagged with FLAG epitope tag to confirm expression from plasmid. Pgk1 was used as loading control. **(e)** Spot analysis of wild-type and *nuc1Δ ski3Δ* strains with plasmid expressing either wild-type *NUC1* or its variants. Strains were grown at the optimal conditions (glucose at 30°C) and the suboptimal conditions (glycerol at 37°C).

We next manipulated Nuc1 mitochondrial localization with heterologous N-terminal targeting sequences attached to a Nuc1-Δ43 construct. The Tom70 signal anchor domain fully restored Nuc1 levels and antiviral activity (Fig 2c-e). Cyb2 is targeted to the mitochondrial intermembrane space through a cleavable bipartite “stop-transfer” mechanism and this targeting sequence similarly restored Nuc1 levels and antiviral activity (Fig 2c-e)^42^. The Cyb2Δ19 variant removes key residues resulting in matrix targeting^42^. We found that Cyb2Δ19-Nuc1 was reduced in abundance and completely defective for limiting L-A levels or of rescuing the *nuc1*Δ *ski3*Δ growth defect (Fig 2c-e). These findings confirm that Nuc1 performs its antiviral function within the mitochondrial intermembrane space.

### Nuc1 governs a cryptic antiviral function of the Rex1 and Rex2 3’-5’ RNA exonucleases

Rex1 and Rex2 are 3’-5’ RNA exonucleases conserved from bacteria to human known to control 3’ end trimming of tRNA, snRNA, and rRNA processing intermediates^22–28^. Previous findings showed that combining *nuc1*Δ with deletions of *REX1* or *REX2* caused synergistic increases in L-A, and we study these interactions further to elucidate the antiviral mechanism^20^.

While Rex1 localizes to the nucleus, Rex2 has been characterized as a mitochondrial protein through biochemical fractionation studies despite its requirement for processing of nuclear RNA species^43–47^. We found that plasmid expressed Rex2-GFP exhibited dual localization to the mitochondria and nucleus, consistent with recent findings in sporulating cells (Fig 3a)^48^. The REXO2 human homolog of Rex2 is also dual localized, but to the mitochondria and cytoplasm. REXO2 localization is controlled through alternative translational initiation that can skip a N-terminal mitochondrial targeting sequence^49,50^. Inspection of *REX2* coding sequences revealed an N-terminal mitochondrial targeting sequence (MTS) with the second AUG codon appearing not until beyond the MTS (M41), exactly as with REXO2 (Fig 3b). Unlike REXO2, Rex2 also contains a C-terminal nuclear localization sequence (Fig 3b). Through mutation of the initiating AUG codon (M1L) or of the M41 encoding AUG codon (M41L) we engineered Rex2-GFP isoforms that exhibited nuclear (M1L) or mitochondrial (M41L) localization (Fig 3a and b). Surprisingly, both mitochondrial and nuclear isoforms of Rex2 restored L-A repression when expressed in a *nuc1*Δ *rex2*Δ strain, though the nuclear form appeared more effective (Fig 3c-e). Mutation of a key predicted catalytic residue (E61A) abolished this rescuing activity (Fig b-e). As both nuclear and mitochondrial Rex2 isoforms provided antiviral function, a developing model is that they do so by repressing RNA species from both compartments that generically enhance L-A replication.

**Figure 3:**
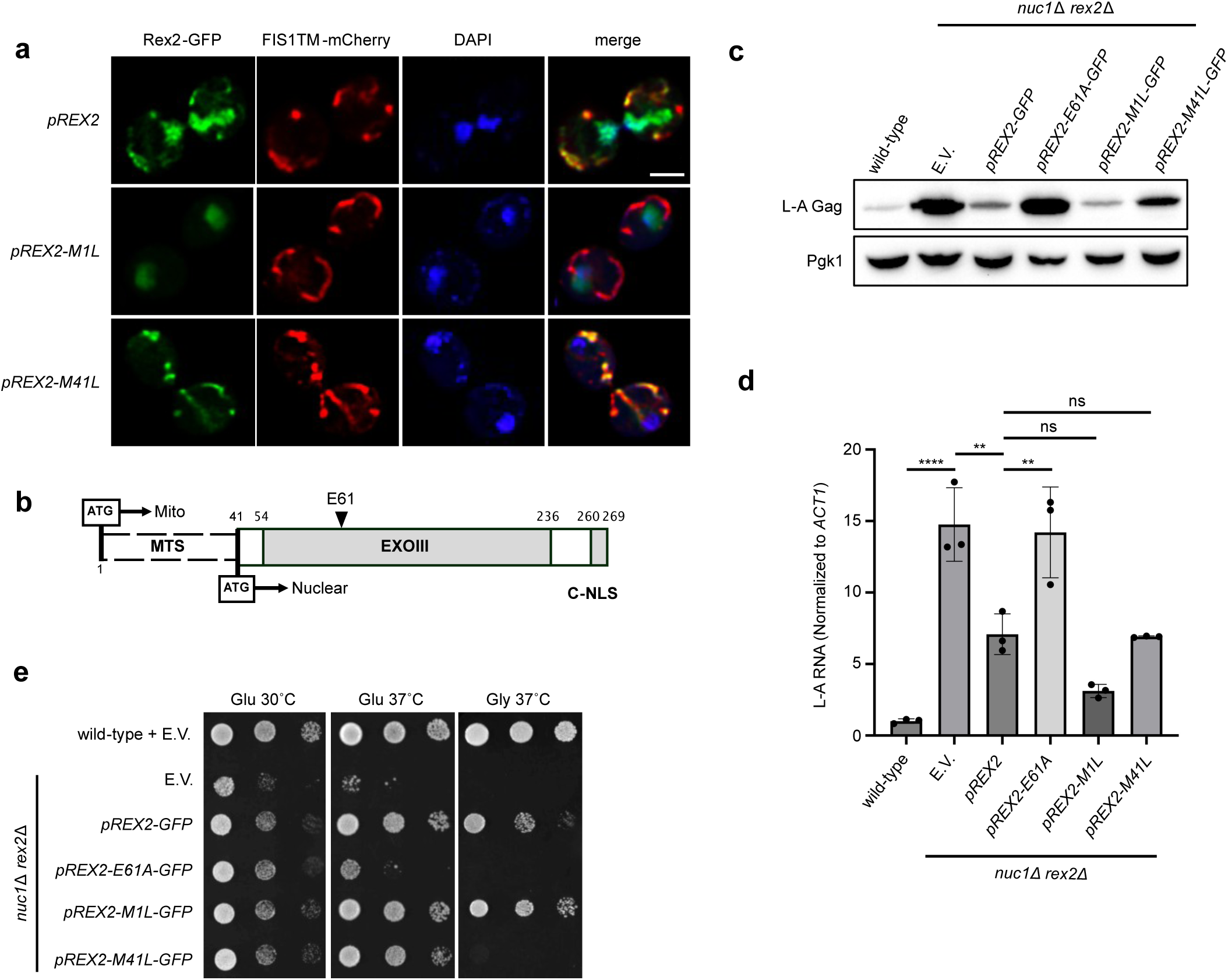
Nuc1 acts synergistically with nuclear and mitochondrial Rex2 antiviral activity. **(a)** Fluorescence microscopy of the subcellular localization of Rex2. Rex2 and its mutated variants were tagged with GFP to monitor its subcellular localization. Mitochondria were marked using an mCherry allele fused to the mitochondrial targeting sequence of Fis1, and cells were stained with DAPI to visualize nuclei. Scale bar, 1 μm. **(b)** Schematic of Rex2 peptide sequence. Regions of interest are highlighted including the two alternative translational initiation start sites, subcellular localization targeting sequence, exonuclease domain, and catalytic residue. **(c)** Western blot of L-A Gag levels in indicated strains. Pgk1 was used as loading control. **(d)** RT-qPCR quantification of L-A RNA normalized to endogenous *ACT1* RNA in indicated strains. Mean RNA level and standard deviation is shown. n = 3. ns = not significant, ** p < 0.01, **** p < 0.0001. The p value was calculated using unpaired student’s t-test. **(e)** Spot analysis of wild-type and *nuc1Δ rex2Δ* strains with plasmid expressing either wild-type *REX2* or its variants. Strains were grown at the optimal conditions (glucose at 30°C) and the increasingly suboptimal conditions (glucose or glycerol at 37°C).

We next measured L-A levels in *nuc1*Δ *rex1*Δ *rex2*Δ triple mutants and observed a massively synergistic interaction resulting in a >200-fold increase compared with wild-type (Fig. 4a). Ethidium staining of electrophoresed total RNA from *nuc1*Δ *rex1*Δ *rex2*Δ triple mutants showed that L-A dsRNA accumulated to levels surpassing those of mature rRNAs, illuminating a remarkable degree of viral replication (Fig. 4b). As expected, *nuc1*Δ *rex1*Δ *rex2*Δ triple mutants exhibited a strong L-A dependent growth defect at high temperature (Fig. 4c). Interestingly, *rex1*Δ *rex2*Δ double mutants did not exhibit any L-A repression defects on their own, nor did they enhance the *ski3*Δ L-A repression defect (Fig 4 a-c). The genetic specificity of these interactions suggest that Nuc1 governs a cryptic antiviral function of *REX1*/*2*.

**Figure 4:**
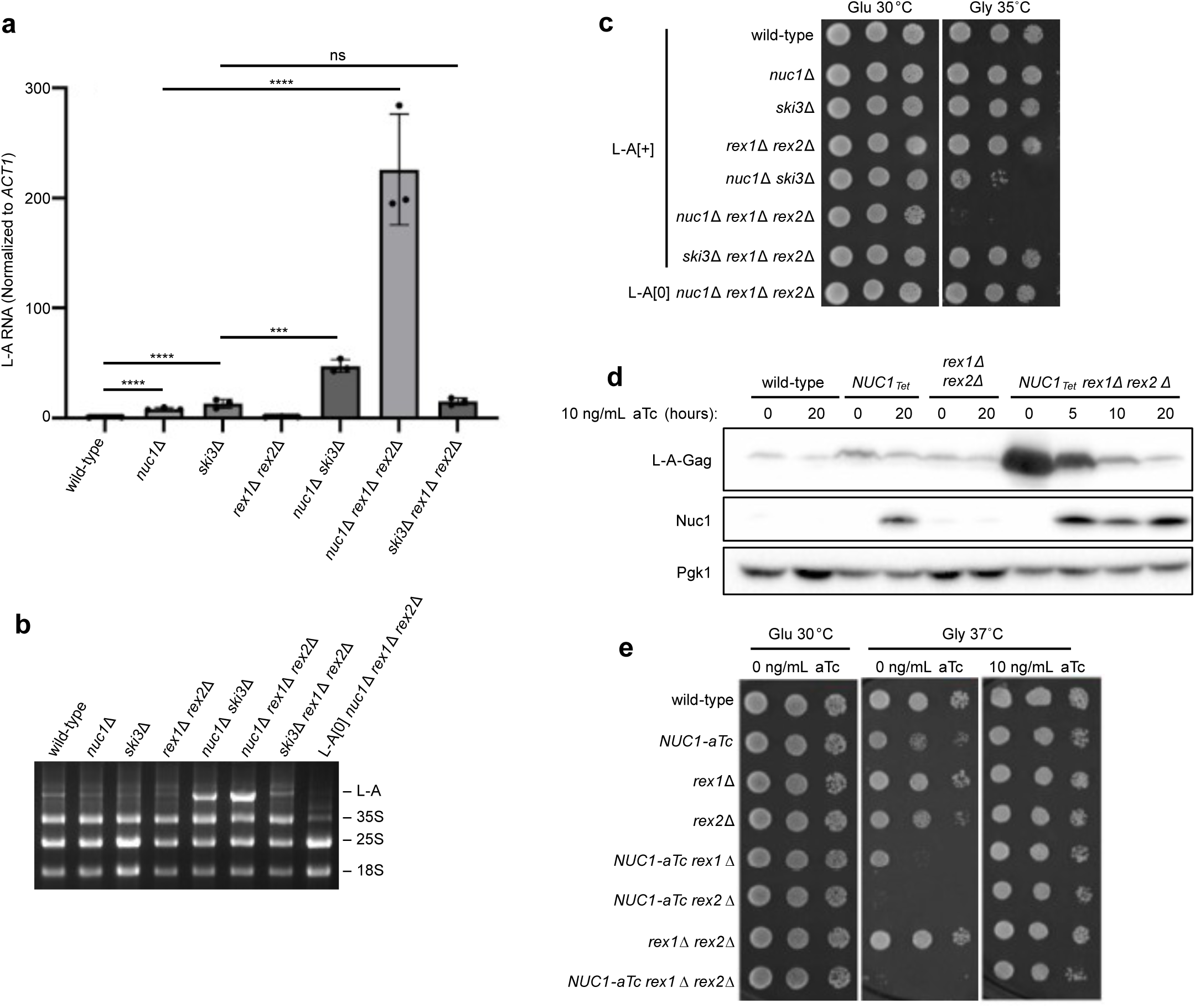
Nuc1 governs the antiviral activity of Rex1 and Rex2. **(a)** RT-qPCR quantification of L-A RNA normalized to endogenous *ACT1* RNA in indicated strains. Mean RNA level and standard deviation is shown. n = 3. ns = not significant, *** p < 0.001, **** p < 0.0001. The p value was calculated using unpaired student’s t-test. **(b)** An ethidium bromide-stained gel of total RNA prepared from indicated strains. The 4.6 kb L-A dsRNA and ribosomal RNAs are indicated. **(c)** Spot analysis of indicated strains. L-A infected strains are denoted as L-A[+] while L-A cured strain is indicated with L-A[0]. Strains were grown at the optimal conditions (glucose at 30°C) and the suboptimal conditions (glycerol at 35°C). **(d)** Western blot of L-A Gag levels in indicated strains. Nuc1 expression was induced by the addition of 10ng/ mL aTc and confirmed using a custom antibody specifically designed to detect Nuc1. Pgk1 was used as loading control. **(e)** Spot analysis of indicated strains. Nuc1 expression was induced by the addition of 10ng/mL aTc. Strains were grown at the optimal conditions (glucose at 30°C) and the suboptimal conditions (glycerol at 37°C).

To further test this model, we engineered a conditional *NUC1* allele using an anhydrotetracycline (aTc) titratable expression system that enables tight repression in the absence of aTc (*NUC1-aTc*)^51^. Under repressing conditions, a *NUC1-aTc rex1*Δ *rex2*Δ strain accumulated massive amounts of L-A Gag leading to growth defects like those of *nuc1*Δ *rex1*Δ *rex2*Δ triple mutants (Fig 4 d, e). Following aTc treatment of this strain Nuc1 protein was induced, Gag levels were restored to wild-type levels over 20 hours of culture (approximately 12 divisions), and the growth defect was rescued (Fig 4 d, e). We conclude that Nuc1 governs a cryptic antiviral function of Rex1 and Rex2 and propose that it does so by preventing aberrant RNA species that accumulate in *rex1*Δ *rex2*Δ from enhancing L-A replication (Fig 5a).

**Figure 5:**
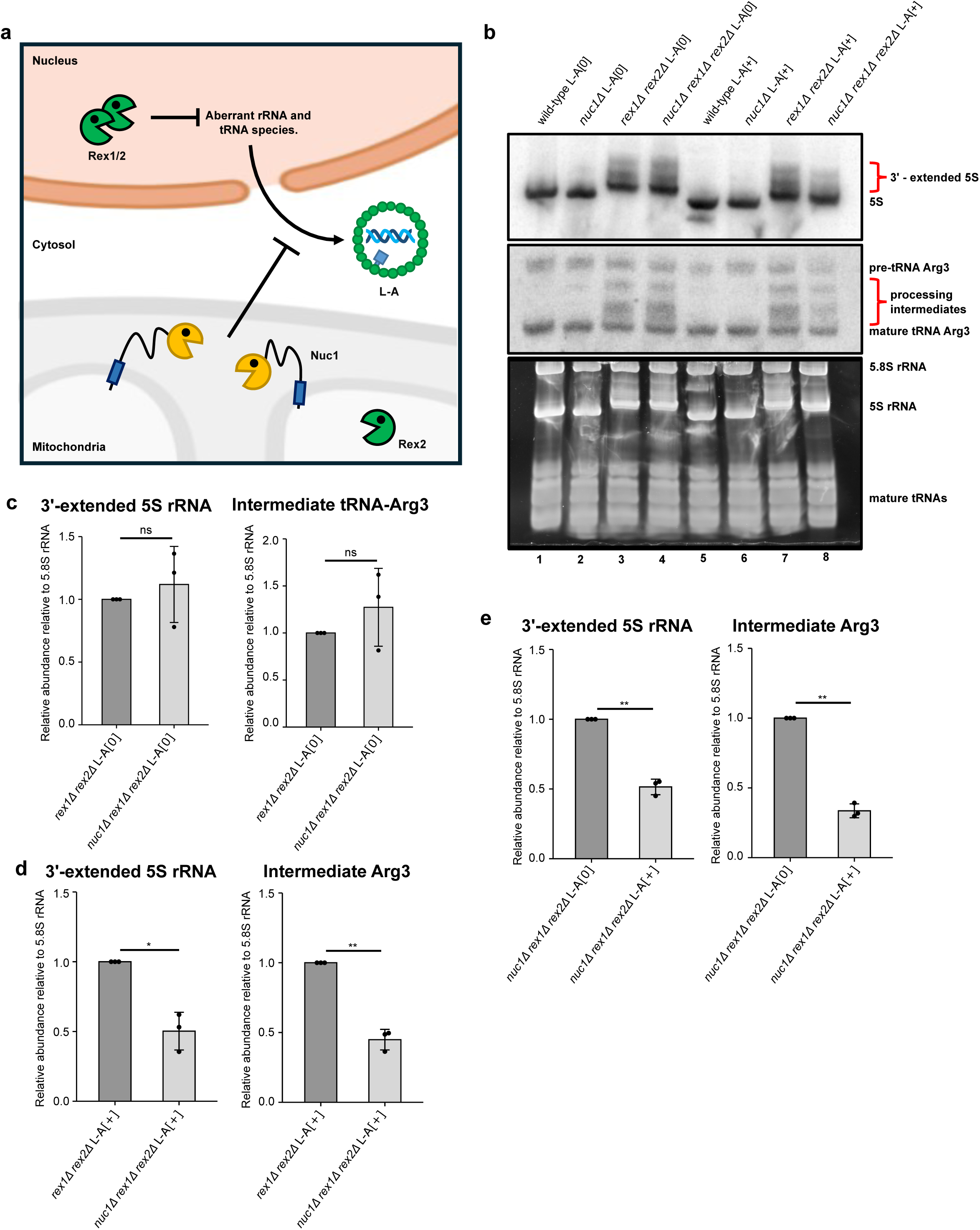
Nuc1 prevents mis-processed 5S rRNA and tRNA-Arg3 from being depleted by L-A. **(a)** Model of Nuc1 antiviral mechanism through depletion of pro-viral non-coding RNA intermediates. Mitochondrial Nuc1 opposes the accumulation of cryptic rRNA and tRNA processing intermediates in *rex1Δ rex2Δ* strains. In the absence of Nuc1 activity, these cryptic RNA intermediates are used to enhance viral replication. **(b)** Northern blot of 5S rRNA and Arg tRNA in indicated strains. L-A infected strains are denoted as L-A[+] while L-A cured strain is indicated with L-A[0]. Ethidium bromide-stained gel of total tRNA and 5.8S rRNA were used as loading control. **(c‒e)** Quantification of 5S rRNA and Arg tRNA processing intermediates from Northern blot normalized to 5.8S rRNA. Relative mean fold change and standard deviation is shown. L-A infected strains are denoted as L-A[+] while L-A cured strain is indicated with L-A[0]. n = 3. * p < 0.05, ** p < 0.01. The p value was calculated using unpaired student’s t-test.

### Nuc1 opposes a pro-viral pool of aberrant rRNA and tRNA processing intermediates

Mutation of *REX1*/*2* causes the accumulation of aberrant processing intermediates of 5S rRNA and tRNA-Arg3^22^. In our model, Nuc1 opposes pro-viral RNAs that accumulate in *rex1*Δ *rex2*Δ mutants, and these known Rex1/2 targets are thus the best candidates to molecularly test it (Fig. 5a). We performed Northern blotting for 5S rRNA and tRNA-Arg3 in wild-type, *nuc1*Δ, *rex1*Δ *rex2*Δ, and *nuc1*Δ *rex1*Δ *rex2*Δ strains that were either uninfected or infected with L-A. These experiments were performed in triplicate using RNA prepared from independent cultures to enable statistical analysis of our findings. A representative replicate is shown (Fig. 5b).

As expected, all *rex1*Δ *rex2*Δ strains accumulated tRNA-Arg3 processing intermediates as well as slower migrating forms of 5S rRNA indicative of known 3’ extensions (Fig. 5b)^22^. The simplest prediction of our model is that these RNAs should increase in abundance if *NUC1* is deleted. While quantifications of the aberrant 5S and tRNA-Arg3 species revealed that they were increased in uninfected *nuc1*Δ *rex1*Δ *rex2*Δ compared with uninfected *rex1*Δ *rex2*Δ, none of these differences attained statistical significance (Fig. 5b, compare red bracketed areas of lanes 3 and 4, and 5c). Curiously, in L-A infected *rex1*Δ *rex2*Δ strains, deletion of *NUC1* led to a statistically significant >2-fold reduced abundance of these aberrant 5S and tRNA-Arg3 species (Fig. 5b, compare red bracketed areas of lanes 7 and 8, and 5d). These results suggest that Nuc1 prevents aberrant 5S and tRNA-Arg3 RNAs from being depleted by the L-A virus. Supporting this interpretation, the depletion of these RNA species in *nuc1*Δ *rex1*Δ *rex2*Δ strains was dependent on L-A infection (Fig. 5b, compare red bracketed areas of lanes 4 and 8, and 5e). Our findings reveal a cryptic RNA surveillance function of Nuc1 that prevents aberrant 5S and tRNA-Arg3 RNAs from being available for consumption by the L-A virus (Fig. 5a). Their fleeting detection in L-A–uninfected *nuc1*Δ *rex1*Δ *rex2*Δ cells suggest that, in the absence of both Nuc1 and L-A, these RNAs are likely subject to further metabolic turnover.

## Discussion

Here we show that Nuc1 governs a novel RNA surveillance system that prevents hyper-replication of the L-A RNA mycovirus. Nuc1 performs this function in the mitochondrial intermembrane space in juxtaposition to L-A’s known cytosolic residence, revealing an indirect mechanism. We elucidate that Nuc1 represses a cryptic pool of aberrantly processed RNAs that accumulate in *rex1*Δ *rex2*Δ mutants These RNAs are depleted by L-A when *NUC1* is mutated. Put simply, our model proposes that aberrant RNAs enter the mitochondria and are subsequently targeted by Nuc1. If Nuc1 activity is not present in the mitochondrial inter-membrane space, they are then available to promote L-A replication, presumably following their export (Fig. 5a).

The movement of RNA into and out of the mitochondria is inherent to our model. Mitochondria are well known targets of extensive protein import, necessitating diverse surveillance systems to maintain proteostasis. One such system imports misfolded cytosolic proteins into mitochondria for degradation^52^. Our model proposes an analogous mitochondrial surveillance system for aberrant 5S rRNA and tRNA-Arg3, in which they are imported into mitochondria for degradation by Nuc1. Other RNAs likely share this fate. Although RNA trafficking between the mitochondria and cytosol remains understudied, numerous noncoding RNAs, including 5S rRNA, various tRNAs, and several catalytic RNAs, are known to be imported into mitochondria in human cells or yeast^53–55^. More recently, double-stranded RNAs have been shown to be exported from mitochondria to the cytosol in human cell lines, triggering a host antiviral response^56^. These findings establish precedent for RNA trafficking both into and out of mitochondria, a key premise of our model.

Another key question our findings raise concerns the mechanism by which *NUC1*-repressed aberrant RNA processing intermediates promote L-A replication. A parsimonious model that explains their L-A dependent depletion is that these RNA species are somehow consumed by L-A, perhaps providing the NTPs necessary to fuel the massive degree of replication we observe in *nuc1*Δ *rex1*Δ *rex2*Δ mutants. It is noteworthy that L-A replicates in discrete viral RNA foci characteristic of “viral factories” that are widely observed in all domains of life. Viral factories compartmentalize the replication, transcription, and chemical modification of viral RNAs, and are the site where diverse host RNA regulatory systems are utilized or even subverted^57^. NTP availability in these factories may be enhanced through scavenging host RNAs that would otherwise be degraded by Nuc1.

Our findings reveal a previously unrecognized form of yeast innate immunity in which Nuc1 depletes aberrant ‘junk’ RNAs from the pool of RNA species available for viral exploitation. This surveillance system may not be dedicated to viral repression *per se*; rather, Nuc1 may serve more broadly as a failsafe mechanism for RNA decay. Whatever the case may be, the responsiveness of L-A to it has provided a sort of molecular magnifying glass, allowing the detection of an otherwise cryptic pool of RNAs controlled by Nuc1. These findings motivate further studies in fungi, bacteria, and metazoans to investigate this new EndoG function.

## Methods

### Yeast strains, media, and plasmids

All strains are isogenic derivatives of BY4742 and were constructed through crossing and dissection as described previously^19,20^. For construction of the anhydrotetracycline regulated allele of *NUC1* (*NUC1*-*aTc*), we followed the methods previously outlined using plasmids provided by Joerg Stelling^51^. Refer to Table S1 for all yeast strains used. All experiments were carried at 30 °C in either YPAD (1% yeast extract, 2% peptone, 0.004% adenine, and 2% glucose) or synthetic complete media (SC; 0.348% yeast nitrogen base, 1% ammonium sulphate, and 2% glucose) with the appropriate amino acid powder mix (Sunrise Science) unless otherwise specified. To induce expression of *NUC1-aTc*, 10ng/mL of anhydrotetracycline was added to the media.

Refer to Table S2 for the plasmids used in this study. The p5476 *2μ LEU2*, p5476 *2μ NIC1 LEU2*, and p5476 *2μ TRZ1 LEU2* were obtained from Brenda Andrews and Charlie Boone^58^. All other plasmids were constructed using the pRS316 and pESC-HIS plasmid backbones and yeast *in vivo* recombination. Mutations were generated using a site-directed mutagenesis kit (NEB; E0554S). For Nuc1 constructs with alternative targeting sequence, PCR fragments of the N-terminus target protein (Tom70 and Cyb2) and Nuc1(Δ43) were amplified from genomic DNA with 21bp overlap on adjacent DNA for assembly purpose. The linearized purified plasmid and gel purified PCR fragments were assembled with NEBuilder HiFi DNA Assembly kit (NEB; E5520S). All plasmids were sequence verified.

### Fluorescent microscopy

Fluorescent microscopy was performed using previously described methods^19,20^. Briefly, cells were fixed with 4% formaldehyde and washed 3 times with 1x phosphate-buffered saline (PBS). Fixed cells were then mounted on a microscope slide and stained with DAPI. Images were acquired with the Leica Sp8 confocal LSM and processed with the Leica Application Suite (LAS X) software package. Deconvolution was applied to all images.

### Yeast subcellular fractionation

The Mitochondrial Yeast Isolation kit and protocol (ab178779; Abcam) was used to fractionate yeast by differential centrifugation as previously described^19^. Briefly, after pelleting the mitochondrial fraction, the soluble cytosolic fraction was precipitated with 2% tricholoroacetic acid at 4 °C for 30 min and washed 3 times with chilled acetone. Both mitochondrial and cytosolic fractions along with whole-cell pellets were resuspended in SDS/PAGE sample buffer (50 mM Tris·HCl, pH 6.8, 2% SDS, 10% glycerol, 0.025% bromophenol blue, and 100 mM dithiothreitol) and heating at 100 °C for 10 min.

### Protein extraction and western blotting

Log-phase cells were harvested and permeabilized with 0.1N NaOH at room temperature for 5 min. The cells were then pelleted and resuspended in SDS/PAGE buffer before heating at 100°C for 10 minutes. The samples were centrifuged to isolate the soluble fraction for western blotting. Protein concentrations were determined with an RC/DC assay (BioRad 5000121). Equal amounts of protein were used for electrophoresis for all samples except for the Nuc1 subcellular fractionation experiment in which ¼ amounts of protein for wild type mitochondrial and total fractions were loaded instead.

Samples were electrophoresed on 10% SDS-PAGE gels and transferred to polyvinylidene difluoride (PVDF) membranes. Membranes were incubated in primary antibody at 4°C overnight and probed with 1:3,000 horseradish peroxidase (HRP)-conjugated horse anti-mouse (7075; Cell Signaling Technology) or goat anti-rabbit (7074; Cell Signaling Technology) secondary antibody. The proteins were detected with Luminata Forte Western HRP Substrate (EMD Millipore) and imaged with the Bio-Rad ChemiDoc XRS+ system. Images were processed with the Image Lab software package (Bio-Rad). The primary antibodies and their dilutions were 1:1,000 anti-FLAG M2 (F1804; Sigma-Aldrich), 1:1,000 anti-Cox2 (obtained from Tom Fox, Cornell University, Ithaca, NY), 1:2,500 anti-GAPDH (obtained from Cordula Enenkel, University of Toronto, Toronto, ON, Canada), 1:5,000 anti-Pgk1 (ab113687; Abcam), and 1:2,000 anti–L-A Gag (obtained from Reed Wickner). A custom polyclonal anti-Nuc1 antibody was generated by CUSABIO (www.cusabio.com) using the full recombinant protein as an immunogen. This was used at a 1:1,000 dilution for the results shown in figure 4d.

### Spot analysis

Yeast strains were grown overnight at 30°C to saturation. Each strain was diluted to an OD600 = 0.4, serial diluted two times by 10-fold and 5 uL was spotted onto agar plates containing synthetic complete media supplemented with 2% glucose, 3% glycerol, or 2% galactose, and grown at 30° C, 35° C, or 37° C as indicated.

### RT-qPCR

RNA was prepared from 10 ODs of cells harvested from log-phase cultures and used for RT-qPCR as previously described^20^. Samples were then incubated for 30 minutes at 65°C in acidic phenol (P4682; Sigma-Aldrich), SDS, and buffer AE (10mM Tris-HCl, 0.5mM EDTA pH 9.0) solution. The phase-separated supernatant was washed with chloroform and precipitated overnight. The precipitate was then washed in 70% ethanol and dissolved in water. RNA samples were purified with the RNeasy Mini Kit (74104; Qiagen), and residual DNA was digested with the RNase-free DNA Removal Kit (AM1906; Invitrogen). RT-qPCR was performed using 1 ng of DNase-treated RNA with the Luna Universal One-Step RT-qPCR Kit (E3005; New England Biolabs). Reactions were prepared according to the manufacturer’s recommendations, with primers at a final concentration of 0.4µM each. Reverse transcription was performed at 55°C for 10 min, followed by initial denaturation at 95°C for 1 min and 40 cycles of 95°C for 10 s and 60°C for 30 s, followed by melt-curve analysis. RT-qPCR was performed on the CFX384 Real-Time PCR Detection System (Bio-Rad). Data were analyzed and plotted using GraphPad Prism.

### Northern blotting

RNA from log phase cultures was prepared as described above. 10μL of 1.4μg/μL RNA was mixed with 2X formamide solution (80% deionized formamide, 0.06% (w/v) bromophenol blue, 0.06% (w/v) xylene cyanol, 10 mM EDTA pH 8.0) in 1:1 ratio. Samples were heated for 5 min at 95°C and snap cooled on ice before being electrophoresed on a 7M urea 10% polyacrylamide gel at 100V at 4°C that had been pre-run for 15 min. Gels were stained in 1X TBE with EtBr and transferred to a positively charged nylon membrane (Revvity - GeneScreen Plus; NEF1017001PK) using an Invitrogen iBlot2 system on setting P0. RNA on the membranes was crosslinked for 1.5 min with a Fotodyne DNA Transfer Lamp (Model 2-1500) and dried for 15 minutes at 80°C using a Model 583 Bio-Rad gel dryer on cycle 2.

Membranes were rotated for 1 hour in 20mL of hybridization solution (6X Saline Sodium Citrate (SSC), 1% Sodium Dodecyl Sulfate (SDS), 2X Denhardt’s solution (Invitrogen; 750018), and 0.1mg/mL yeast RNA) at 10°C lower than the corresponding probe melting temperature (Tm). Refer to Table S3 for the probe sequences. 2.5⌠M DNA probes were 5’ γ-^32^P-ATP labelled with T4 Polynucleotide Kinase (NEB; M0201S) in a 10⌠L reaction for 2h at 37°C, heated at 95°C for 5 min, and added to the membrane hybridization solution for rotation overnight at Tm -10°C. Hybridization solution was removed, and membranes were rotated 3X for 20 min with 30mL wash buffer (2X SSC, 0.1% SDS) at Tm -10°C, Tm - 20°C, and Tm -30°C respectively prior to overnight exposure to a Phosphor Screen, and phosphor-imaging using a Typhoon. Between probe hybridizations, membranes were stripped by rotation 3X for 20 min with 30mL strip buffer (0.1X SSC, 0.1% SDS) at 70°C.

RNA signal was calculated using Cytiva ImageQuant Software and normalized by subtracting each RNA volumes from probe background volume and dividing by the corresponding 5.8S EtBr volume in each lane. Relative abundance of each RNA species was calculated by setting baseline to 1.0 and completing a two-tailed paired t-test for each direct comparison.

---

**Figure S1:**
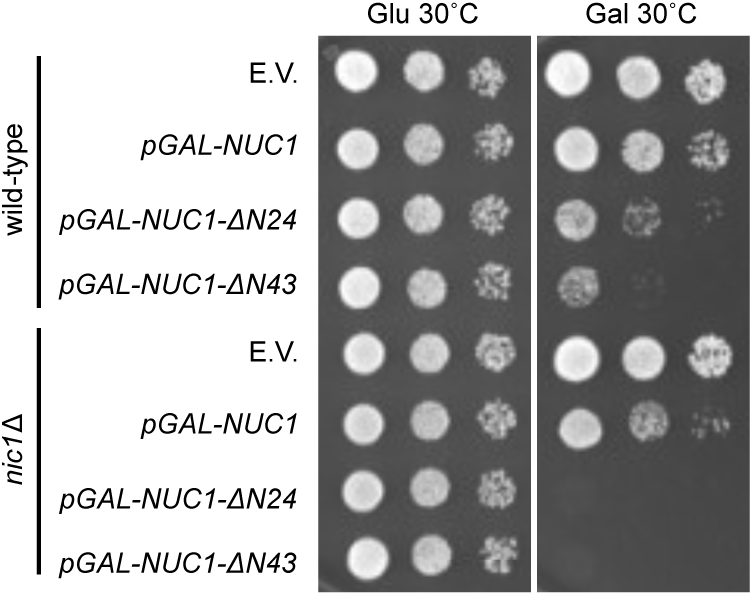
Nuc1’s cytosolic localization is toxic to the cells. Spot analysis of wild-type and *nic1Δ* strains with plasmid expressing either wild-type *NUC1* or N-terminus deleted *NUC1* alleles. *NUC1* overexpression was induced by galactose treatment.

**Table S1:**
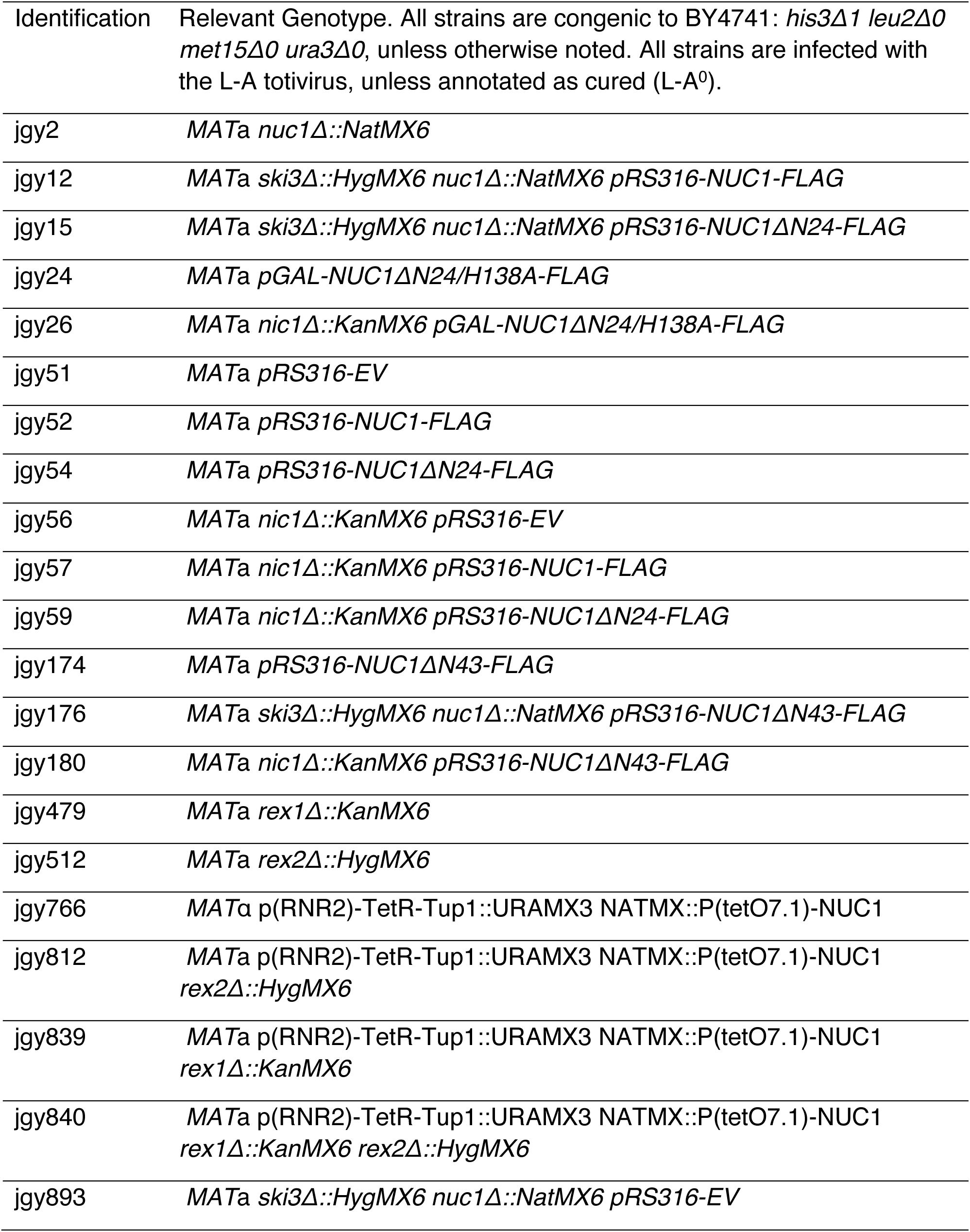
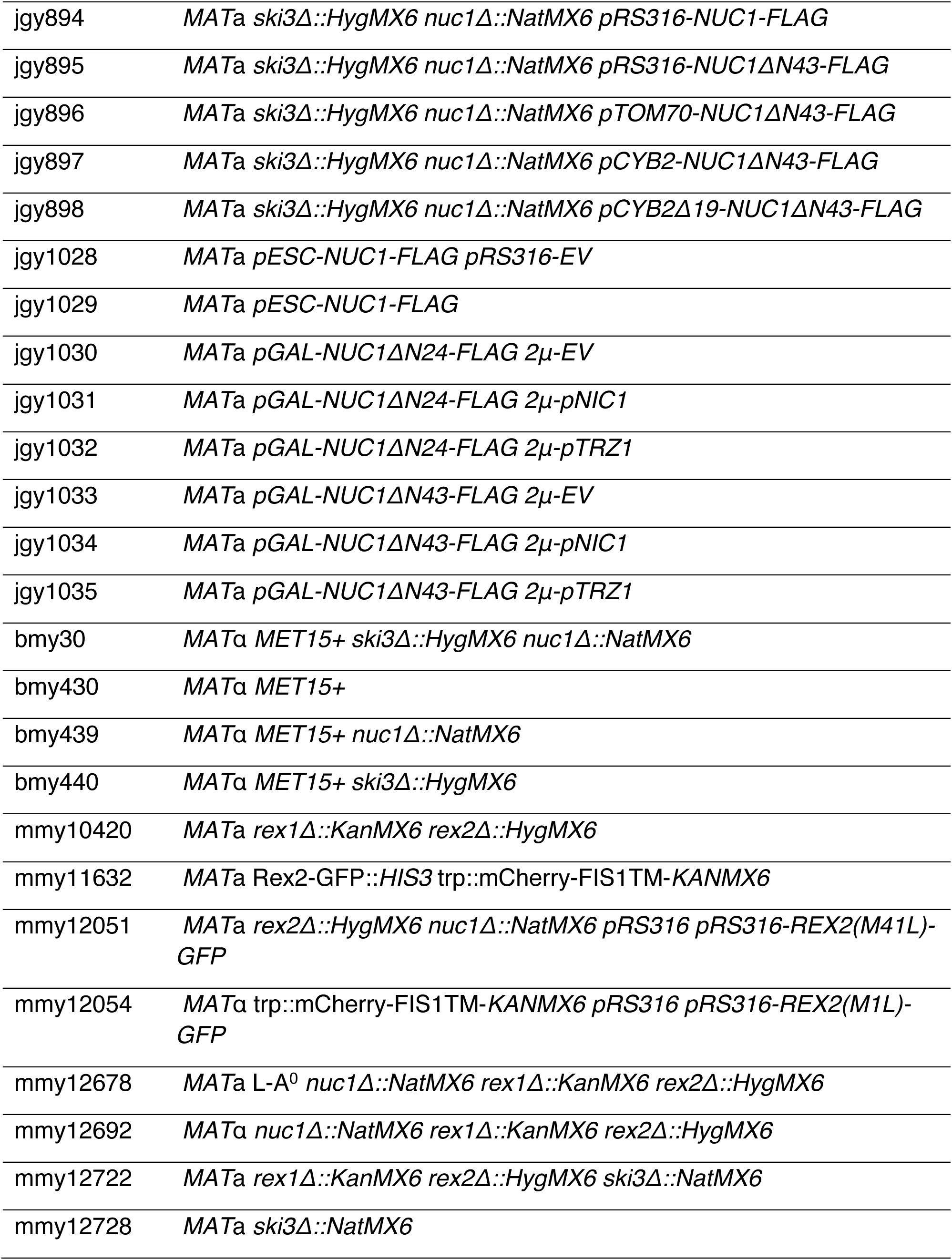
Strains.

**Table S2:** Plasmids.

| Identifier | Plasmid construct |
| --- | --- |
| mmb92 | pRS316 CEN URA3 AMP |
| mmb235 | pESC-GAL1/10pr::*NUC1*-FLAG 2μ HIS3 AMP |
| mmb332 | p5476 2μ LEU2 AMP |
| mmb841 | pRS316 *NUC1*-FLAG |
| mmb844 | p5476 p*NIC1* |
| mmb869 | pRS316 *NUC1*(ΔN24)-FLAG |
| mmb870 | pRS316 *NUC1*(ΔN24, H138A)-FLAG |
| mmb871 | pESC GAL1/10pr::*NUC1*(ΔN24)-FLAG |
| mmb872 | pESC GAL1/10pr::*NUC1*(ΔN24, H138A)-FLAG |
| mmb920 | pESC GAL1/10pr::*NUC1*(ΔN43)-FLAG |
| mmb997 | pRS316 *NUC1*(ΔN43)-FLAG |
| mmb1113 | pRS316 *REX2*-GFP |
| mmb1145 | pRS316 *REX2*(E61A)-GFP |
| mmb1146 | pRS316 *REX2*(M1L)-GFP |
| mmb1147 | pRS316 *REX2*(M41L)-GFP |
| mmb1219 | pRS316 *TOM70*-*NUC1*(ΔN43)-FLAG |
| mmb1220 | pRS316 *CYB2*-*NUC1*(ΔN43)-FLAG |
| mmb1221 | pRS316 *CYB2*(Δ19)-*NUC1*(ΔN43)-FLAG |
| mmb1343 | p5476 p*TRZ1* |

**Table S3:**
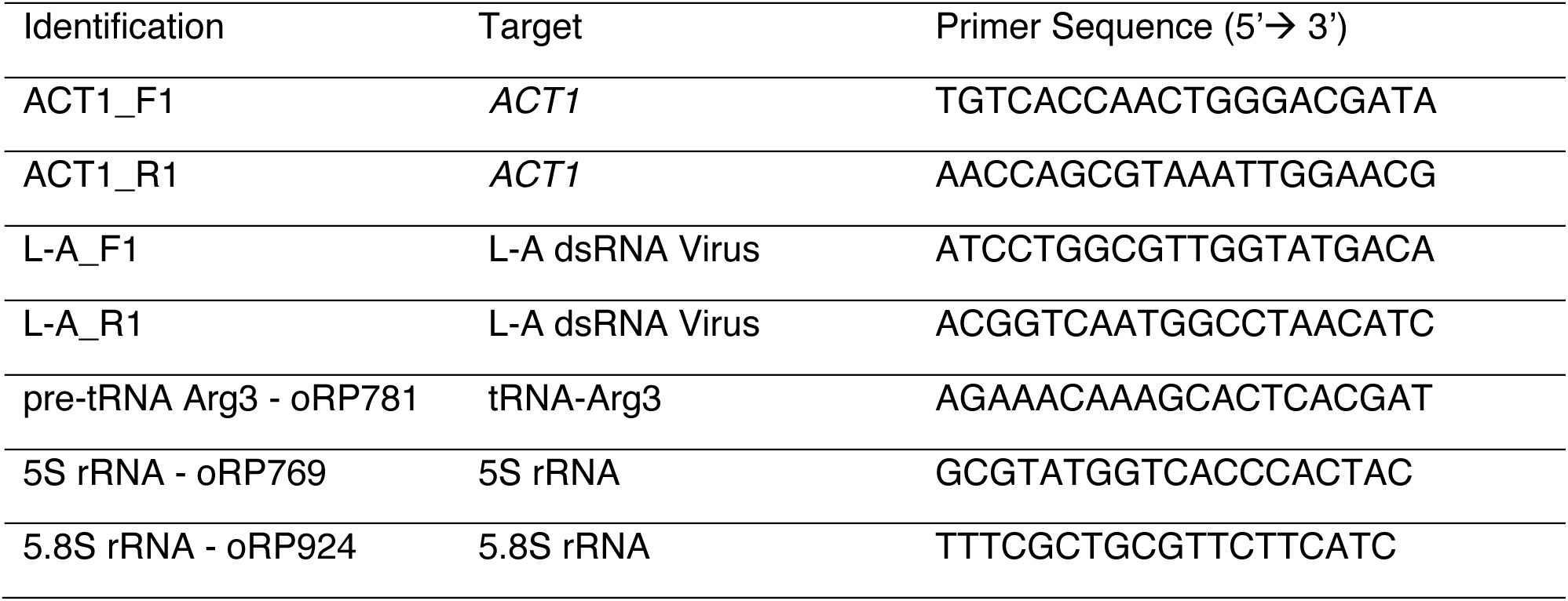
Primers.

## References

1 Cymerman, I. A., Chung, I., Beckmann, B. M., Bujnicki, J. M. & Meiss, G. EXOG, a novel paralog of Endonuclease G in higher eukaryotes. Nucleic Acids Res 36, 1369–1379, doi:10.1093/nar/gkm1169 (2008).

2 Derre-Bobillot, A. et al. Nuclease A (Gbs0661), an extracellular nuclease of Streptococcus agalactiae, attacks the neutrophil extracellular traps and is needed for full virulence. Mol Microbiol 89, 518–531, doi:10.1111/mmi.12295 (2013).

3 Liao, C., Mao, F., Qian, M. & Wang, X. Pathogen-Derived Nucleases: An Effective Weapon for Escaping Extracellular Traps. Front Immunol 13, 899890, doi:10.3389/fimmu.2022.899890 (2022).

4 Zhang, Y. et al. A Conserved Endonuclease From Rust Fungi Suppresses Plant Extracellular DNA-Triggered Immunity. Plant Cell Environ 48, 4356–4369, doi:10.1111/pce.15440 (2025).

5 Li, L. Y., Luo, X. & Wang, X. Endonuclease G is an apoptotic DNase when released from mitochondria. Nature 412, 95–99, doi:10.1038/35083620 (2001).

6 Zhou, Q. et al. Mitochondrial endonuclease G mediates breakdown of paternal mitochondria upon fertilization. Science 353, 394–399, doi:10.1126/science.aaf4777 (2016).

7 DeLuca, S. Z. & O’Farrell, P. H. Barriers to male transmission of mitochondrial DNA in sperm development. Dev Cell 22, 660–668, doi:10.1016/j.devcel.2011.12.021 (2012).

8 Kim, J. et al. VDAC oligomers form mitochondrial pores to release mtDNA fragments and promote lupus-like disease. Science 366, 1531–1536, doi:10.1126/science.aav4011 (2019).

9 Dake, E., Hofmann, T. J., McIntire, S., Hudson, A. & Zassenhaus, H. P. Purification and properties of the major nuclease from mitochondria of Saccharomyces cerevisiae. J Biol Chem 263, 7691–7702 (1988).

10 Cote, J. & Ruiz-Carrillo, A. Primers for mitochondrial DNA replication generated by endonuclease G. Science 261, 765–769, doi:10.1126/science.7688144 (1993).

11 Kalinowska, M., Garncarz, W., Pietrowska, M., Garrard, W. T. & Widlak, P. Regulation of the human apoptotic DNase/RNase endonuclease G: involvement of Hsp70 and ATP. Apoptosis 10, 821–830, doi:10.1007/s10495-005-0410-9 (2005).

12 Xu, X. et al. Endonuclease G promotes hepatic mitochondrial respiration by selectively increasing mitochondrial tRNA(Thr) production. Proc Natl Acad Sci U S A 122, e2411298122, doi:10.1073/pnas.2411298122 (2025).

13 Nakashima, A. et al. Genes for a nuclease and a protease are involved in the drastic decrease in cellular RNA amount in fission yeast cells during nitrogen starvation. J Biochem 131, 391–398, doi:10.1093/oxfordjournals.jbchem.a003114 (2002).

14 Yu, Y. et al. Yeast EndoG prevents genome instability by degrading extranuclear DNA species. Nat Commun 15, 7653, doi:10.1038/s41467-024-52147-2 (2024).

15 Cupo, R. R., Dominguez-Martin, E. & Youle, R. The mitochondrial intermembrane space nuclease Nuc1 (endonuclease G) prevents mitophagy-mediated mtDNA escape in yeast. Curr Biol 36, 912–927 e914, doi:10.1016/j.cub.2026.01.006 (2026).

16 Buttner, S. et al. Endonuclease G regulates budding yeast life and death. Mol Cell 25, 233–246, doi:10.1016/j.molcel.2006.12.021 (2007).

17 Eastwood, M. D., Cheung, S. W., Lee, K. Y., Moffat, J. & Meneghini, M. D. Developmentally programmed nuclear destruction during yeast gametogenesis. Dev Cell 23, 35–44, doi:10.1016/j.devcel.2012.05.005 (2012).

18 Liu, Y. X. & Dieckmann, C. L. Overproduction of yeast viruslike particles by strains deficient in a mitochondrial nuclease. Mol Cell Biol 9, 3323–3331, doi:10.1128/mcb.9.8.3323-3331.1989 (1989).

19 Gao, J. et al. Meiotic viral attenuation through an ancestral apoptotic pathway. Proc Natl Acad Sci U S A 116, 16454–16462, doi:10.1073/pnas.1900751116 (2019).

20 Chau, S. et al. Diverse yeast antiviral systems prevent lethal pathogenesis caused by the L-A mycovirus. Proc Natl Acad Sci U S A 120, e2208695120, doi:10.1073/pnas.2208695120 (2023).

21 Wickner, R. B. & Edskes, H. K. The L-A dsRNA Virus and Its Satellites: Totiviruses and Killers in Saccharomyces cerevisiae. Viruses 18, doi:10.3390/v18080920 (2026).

22 van Hoof, A., Lennertz, P. & Parker, R. Three conserved members of the RNase D family have unique and overlapping functions in the processing of 5S, 5.8S, U4, U5, RNase MRP and RNase P RNAs in yeast. EMBO J 19, 1357–1365, doi:10.1093/emboj/19.6.1357 (2000).

23 Piper, P. W., Bellatin, J. A. & Lockheart, A. Altered maturation of sequences at the 3’ terminus of 5S gene transcripts in a Saccharomyces cerevisiae mutant that lacks a RNA processing endonuclease. EMBO J 2, 353–359, doi:10.1002/j.1460-2075.1983.tb01430.x (1983).

24 Copela, L. A., Fernandez, C. F., Sherrer, R. L. & Wolin, S. L. Competition between the Rex1 exonuclease and the La protein affects both Trf4p-mediated RNA quality control and pre-tRNA maturation. RNA 14, 1214–1227, doi:10.1261/rna.1050408 (2008).

25 Ozanick, S. G. et al. Rex1p deficiency leads to accumulation of precursor initiator tRNAMet and polyadenylation of substrate RNAs in Saccharomyces cerevisiae. Nucleic Acids Res 37, 298–308, doi:10.1093/nar/gkn925 (2009).

26 Skowronek, E., Grzechnik, P., Spath, B., Marchfelder, A. & Kufel, J. tRNA 3’ processing in yeast involves tRNase Z, Rex1, and Rrp6. RNA 20, 115–130, doi:10.1261/rna.041467.113 (2014).

27 Foretek, D., Wu, J., Hopper, A. K. & Boguta, M. Control of Saccharomyces cerevisiae pre-tRNA processing by environmental conditions. RNA 22, 339–349, doi:10.1261/rna.054973.115 (2016).

28 An, W., Yan, Y. & Ye, K. High resolution landscape of ribosomal RNA processing and surveillance. Nucleic Acids Res 52, 10630–10644, doi:10.1093/nar/gkae606 (2024).

29 Waizenegger, T., Stan, T., Neupert, W. & Rapaport, D. Signal-anchor domains of proteins of the outer membrane of mitochondria: structural and functional characteristics. J Biol Chem 278, 42064–42071, doi:10.1074/jbc.M305736200 (2003).

30 Ghosh, M., Meiss, G., Pingoud, A. M., London, R. E. & Pedersen, L. C. The nuclease a-inhibitor complex is characterized by a novel metal ion bridge. J Biol Chem 282, 5682–5690, doi:10.1074/jbc.M605986200 (2007).

31 Muro-Pastor, A. M., Herrero, A. & Flores, E. The nuiA gene from Anabaena sp. encoding an inhibitor of the NucA sugar-non-specific nuclease. J Mol Biol 268, 589–598, doi:10.1006/jmbi.1997.0985 (1997).

32 Temme, C. et al. The Drosophila melanogaster Gene cg4930 Encodes a High Affinity Inhibitor for Endonuclease G. J Biol Chem 284, 8337–8348, doi:10.1074/jbc.M808319200 (2009).

33 Loll, B., Gebhardt, M., Wahle, E. & Meinhart, A. Crystal structure of the EndoG/EndoGI complex: mechanism of EndoG inhibition. Nucleic Acids Res 37, 7312–7320, doi:10.1093/nar/gkp770 (2009).

34 Batsaikhan, J. et al. KU80 suppresses endonuclease G activity to preserve genomic integrity. FEBS J 293, 5952–5966, doi:10.1111/febs.70495 (2026).

35 Graille, M. et al. Structure-based functional annotation: yeast ymr099c codes for a D-hexose-6-phosphate mutarotase. J Biol Chem 281, 30175–30185, doi:10.1074/jbc.M604443200 (2006).

36 Huh, W. K. et al. Global analysis of protein localization in budding yeast. Nature 425, 686–691, doi:10.1038/nature02026 (2003).

37 Ma, M. et al. Trz1, the long form RNase Z from yeast, forms a stable heterohexamer with endonuclease Nuc1 and mutarotase. Biochem J 474, 3599–3613, doi:10.1042/BCJ20170435 (2017).

38 Hazbun, T. R. et al. Assigning function to yeast proteins by integration of technologies. Mol Cell 12, 1353–1365, doi:10.1016/s1097-2765(03)00476-3 (2003).

39 Gavin, A. C. et al. Proteome survey reveals modularity of the yeast cell machinery. Nature 440, 631–636, doi:10.1038/nature04532 (2006).

40 Babu, M. et al. Interaction landscape of membrane-protein complexes in Saccharomyces cerevisiae. Nature 489, 585–589, doi:10.1038/nature11354 (2012).

41 Benschop, J. J. et al. A consensus of core protein complex compositions for Saccharomyces cerevisiae. Mol Cell 38, 916–928, doi:10.1016/j.molcel.2010.06.002 (2010).

42 Beasley, E. M., Muller, S. & Schatz, G. The signal that sorts yeast cytochrome b2 to the mitochondrial intermembrane space contains three distinct functional regions. EMBO J 12, 2303–2311, doi:10.1002/j.1460-2075.1993.tb05884.x (1993).

43 Hanekamp, T. & Thorsness, P. E. YNT20, a bypass suppressor of yme1 yme2, encodes a putative 3’-5’ exonuclease localized in mitochondria of Saccharomyces cerevisiae. Curr Genet 34, 438–448, doi:10.1007/s002940050418 (1999).

44 Sickmann, A. et al. The proteome of Saccharomyces cerevisiae mitochondria. Proc Natl Acad Sci U S A 100, 13207–13212, doi:10.1073/pnas.2135385100 (2003).

45 Reinders, J., Zahedi, R. P., Pfanner, N., Meisinger, C. & Sickmann, A. Toward the complete yeast mitochondrial proteome: multidimensional separation techniques for mitochondrial proteomics. J Proteome Res 5, 1543–1554, doi:10.1021/pr050477f (2006).

46 Renvoise, M. et al. Quantitative variations of the mitochondrial proteome and phosphoproteome during fermentative and respiratory growth in Saccharomyces cerevisiae. J Proteomics 106, 140–150, doi:10.1016/j.jprot.2014.04.022 (2014).

47 Frank, P., Braunshofer-Reiter, C., Karwan, A., Grimm, R. & Wintersberger, U. Purification of Saccharomyces cerevisiae RNase H(70) and identification of the corresponding gene. FEBS Lett 450, 251–256, doi:10.1016/s0014-5793(99)00512-8 (1999).

48 Higdon, A. L., Won, N. H. & Brar, G. A. Truncated protein isoforms generate diversity of protein localization and function in yeast. Cell Syst 15, 388–408 e384, doi:10.1016/j.cels.2024.03.005 (2024).

49 Szewczyk, M. et al. Human REXO2 controls short mitochondrial RNAs generated by mtRNA processing and decay machinery to prevent accumulation of double-stranded RNA. Nucleic Acids Res 48, 5572–5590, doi:10.1093/nar/gkaa302 (2020).

50 Bruni, F., Gramegna, P., Oliveira, J. M., Lightowlers, R. N. & Chrzanowska-Lightowlers, Z. M. REXO2 is an oligoribonuclease active in human mitochondria. PLoS One 8, e64670, doi:10.1371/journal.pone.0064670 (2013).

51 Azizoglu, A., Loureiro, C., Venetz, J. & Brent, R. Autorepression-Based Conditional Gene Expression System in Yeast for Variation-Suppressed Control of Protein Dosage. Curr Protoc 3, e647, doi:10.1002/cpz1.647 (2023).

52 Ruan, L. et al. Cytosolic proteostasis through importing of misfolded proteins into mitochondria. Nature 543, 443–446, doi:10.1038/nature21695 (2017).

53 Sharma, S. & Fazal, F. M. Localization of RNAs to the mitochondria-mechanisms and functions. RNA 30, 597–608, doi:10.1261/rna.079999.124 (2024).

54 Kolesnikova, O. et al. Targeting of tRNA into yeast and human mitochondria: the role of anticodon nucleotides. Mitochondrion 2, 95–107, doi:10.1016/s1567-7249(02)00013-2 (2002).

55 Muneretto, G., Plazzi, F. & Passamonti, M. Mitochondrion-to-nucleus communication mediated by RNA export: a survey of potential mechanisms and players across eukaryotes. Biol Lett 20, 20240147, doi:10.1098/rsbl.2024.0147 (2024).

56 Dhir, A. et al. Mitochondrial double-stranded RNA triggers antiviral signalling in humans. Nature 560, 238–242, doi:10.1038/s41586-018-0363-0 (2018).

57 Brittain, K. S., Delfino, L. M., Thiemann, O. H., Cortines, J. R. & Parent, K. N. The organization and dynamics of viral factories. J Virol 100, e0089226, doi:10.1128/jvi.00892-26 (2026).

58 Ho, C. H. et al. A molecular barcoded yeast ORF library enables mode-of-action analysis of bioactive compounds. Nat Biotechnol 27, 369–377, doi:10.1038/nbt.1534 (2009).

